# Integrative analysis reveals RNA G-Quadruplexes in UTRs are selectively constrained and enriched for functional associations

**DOI:** 10.1101/666842

**Authors:** David S.M. Lee, Louis R. Ghanem, Yoseph Barash

## Abstract

Identifying regulatory elements in the noncoding genome is a fundamental challenge in biology. G-quadruplex (G4) sequences are abundant in untranslated regions (UTRs) of human messenger RNAs, but their functional importance remains unclear. By integrating multiple sources of genetic and genomic data, we show that putative G-quadruplex forming sequences (pG4) in 5’ and 3’ UTRs are selectively constrained, and enriched for cis-eQTLs and RNA-binding protein (RBP) interactions. Using over 15,000 whole-genome sequences, we uncover a degree of negative (purifying) selection in UTR pG4s comparable to that of missense variation in protein-coding sequences. In parallel, we identify new proteins with evidence for preferential binding at pG4s from ENCODE annotations, and delineate putative regulatory networks composed of shared binding targets. Finally, by mapping variants in the NIH GWAS Catalogue and ClinVar, we find enrichment for disease-associated variation in 3’UTR pG4s. At a GWAS pG4-variant associated with hypertension in HSPB7, we uncover robust allelic imbalance in GTEx RNA-seq across multiple tissues, suggesting that changes in gene expression associated with pG4 disruption underlie the observed phenotypic association. Taken together, our results establish UTR G-quadruplexes as important cis-regulatory features, and point to a putative link between disruption within UTR pG4 and susceptibility to human disease.

## INTRODUCTION

Interpreting the impact of genetic variation in noncoding regions of the genome remains challenging. Indeed, most disease-associated genetic variants discovered through genome-wide association studies occupy noncoding regions of the genome, highlighting the importance of understanding how variation in these regions might perturb regulatory mechanisms that contribute to disease pathology^1^. The 5’ and 3’ untranslated regions (UTRs) are included within mature mRNA transcripts and can significantly regulate gene expression via post-transcriptional mechanisms. Thus, understanding the impact of cis-regulatory variants within UTRs is of critical importance towards improving the interpretation of noncoding genetic variation in human health and disease.

Guanine rich nucleic acid sequences can form non-canonical secondary structures known as G-quadruplexes (G4s) in both DNA and RNA^2^. In contrast to DNA G-quadruplexes, RNA G4s form more readily *in vitro* due to their greater thermodynamic stability and reduced steric hindrance ^3,4^. Both bioinformatics and experimental approaches towards identifying RNA G4s have uncovered abundant putative RNA G-quadruplex (pG4) forming regions within the human genome, and it has been observed that G4s are enriched within both the 5’ and 3’ UTRs of messenger RNAs^5^.

The relative abundance of pG4 sequences within mRNA UTRs suggests that they serve functionally important roles in regulating gene expression. Although specific RNA G4s have been experimentally associated with diverse biological functions, including mediating translational control^6,7^, alternative splicing^8^, subcellular localization^9^, and RNA stability^10,11^, the overall functional importance of RNA G4s in the human transcriptome has largely been extrapolated from a limited number of experimental studies, and remains unresolved. This uncertainty over the biological significance of RNA G4s is highlighted by recent findings suggesting that almost all detectable RNA G4s exist as unfolded RNA at steady-state *in cellulo*^12^. Thus the question of whether these non-canonical secondary structures serve important biological functions within the cell remains unresolved.

To address this question, we combined several large scale genomic and genetic data resources to assess evidence for evolutionary constraint on UTR pG4 sequences in humans, and enrichment for functional associations, including cis-eQTLs and protein binding sites. Specifically, we use whole genome sequencing of over 15,000 individuals from gnomAD^13^ to compare the frequency and distribution of single nucleotide variants (SNVs) within pG4 sequences in UTRs compared to coding sequences, and non-pG4-forming UTR regions. We then address the potential functional significance of these UTR pG4s by using GTEx to assess their expression across 53 tissue contexts, and their enrichment for annotated cis-eQTLs. In parallel, we evaluate colocalization between UTR pG4 and protein binding sites observed in cross-linking immunoprecipitation sequencing (CLIP-seq) experiments from ENCODE, and elucidate new modules of post-transcriptional regulation based on shared pG4-gene binding targets. Finally, we show that disease-associated genes in ClinVar are enriched for variation within pG4 sequences compared to non-pG4 UTR sequences; and identify a GWAS variant associated with hypertension within a 3’UTR pG4 in HSPB7 that is also a cis-eQTL, suggesting that increased HSPB7 expression is associated with increased susceptibility to hypertension.

Overall, our results establish UTR pG4 sequences as evolutionarily constrained features within annotated protein-coding mRNAs in the human transcriptome. We demonstrate that these constrained UTR pG4 sequences exhibit enrichment for cis-eQTL variants as identified by GTEx, and find subsets of RNA binding proteins (RBPs) that are selectively enriched within 5’ and 3’ UTR pG4 sequences. Taken together, our results point to the biological significance of RNA pG4 sequences within UTRs, and highlight the importance of considering structure in determining biological function in noncoding regions of the genome.

## RESULTS

### pG4 sequences exhibit heightened selective pressure within UTRs

Given the reported enrichment of G4 sequences within human UTRs, we hypothesized that RNA G4s within these regions should exhibit patterns of genetic variation consistent with heightened evolutionary constraint in humans. To test this hypothesis we first mapped pG4 sequences transcriptome-wide within annotated UTRs. Although diverse sequences have been reported to form *in vitro* G-quadruplexes^14^, we chose to focus on the canonical G4 motif – GGG-{N-1:7}(3)-GGG (**Figure 1a**) to limit the potential number of false-positive G4s in downstream analyses (see methods). Using a text-based pattern matching approach over the Ensemble transcript database (release v. 75), we identified 2967 unique protein-coding genes encoding for at least one transcript isoform containing a pG4 sequence within the 5’UTR, and 2835 protein-coding genes with at least one transcript isoform encoding a pG4 sequence within the 3’UTR (suppl. Table 1). These numbers are consistent with previously reported results from a similar G-quadruplex identification approach^5^. To further increase the specificity of pG4 sequences, we also defined a set of experimentally supported “rG4” sequences (466 in the 5’UTR, 1743 in the 3’UTR), consisting of canonical pG4 sequence with evidence of secondary structure formation in HeLa cells as determined by *in vitro* structure mapping approaches (see **methods**)^15^.

**Figure 1:**
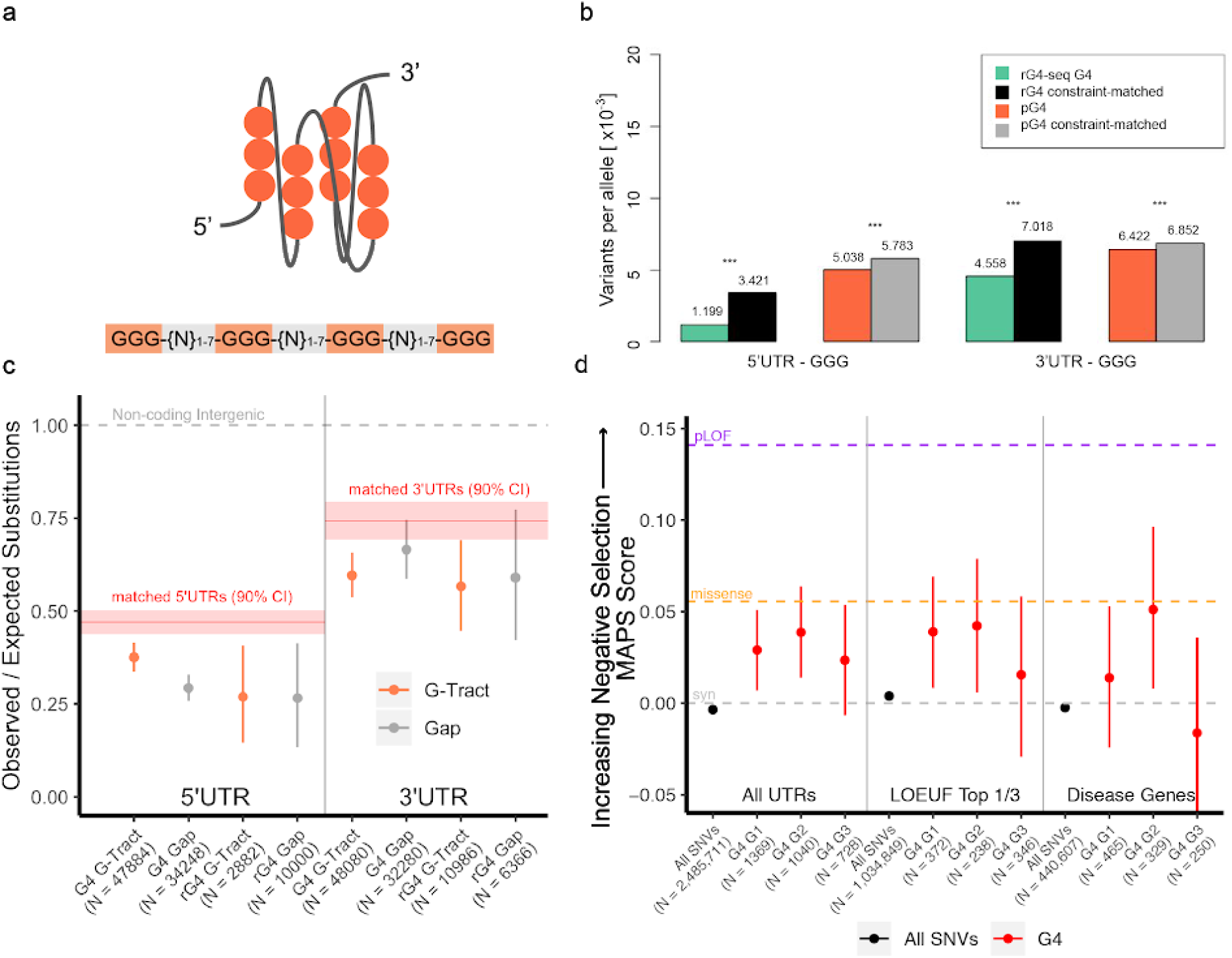
UTR pG4 sequences exhibit greater sequence constraint compared to non-pG4 forming UTR sequences. (a) Schematic depicting a folded RNA parallel G-Quadruplex with the accompanying “canonical” pG4-sequence (GGG-{N:1-7}-GGG{N:1-7}-GGG{N:1-7}-GGG). (b) Reduction in variant frequencies affecting guanine G-tracts within UTR pG4 forming sequences compared to matched non-pG4 G-tracts by transcript-level constraint. *** denotes P-value < 2.2e-16 by Fisher’s Exact Test. (c) Reduction in the number of observed polymorphic sites compared to expectation in 5’ and 3’UTR pG4 forming G-tracts using a nucleotide substitution model based on local sequence context (permuted *P*<1×10^−4^ in all G-tracts compared to match non-pG4 UTR sequences). Bars represent bootstrapped 90% confidence interval for the ratio of observed versus expected substitutions within each pG4 region. “rG4”-G-tracts are those within UTR pG4 that have evidence of secondary structure formation by rG4-seq (see **methods**). Red line and shaded regions represent the observed versus expected number of substitutions in non-pG4 UTR sequences matched by transcript-level constraint and 90% confidence intervals respectively. Grey-dashed line represents an expected versus observed ratio of 1:1. (d) Mutability-Adjusted Proportion of Singletons (MAPS) for each set of set of variants affecting trinucleotide guanines within the “meta” pG4 sequence motif. Central position guanines consistently demonstrate the highest MAPS scores (are most constrained) compared to non-pG4 UTR variants (permuted *P*<1×10^−4^) across all contexts. Bars represent the 5% and 95% bootstrap permutations for each variant class. Purple-dashed line “pLOF”, orange-dashed line, and grey-dashed line represent MAPS score for Ensemble predicted “high-impact” coding, missense, and synonymous mutations respectively.

Next, we assessed relative degrees of sequence constraint across UTR pG4 and non-pG4 forming sequences. Using whole genome sequencing data from over 15,000 individuals from the public gnomAD release (version 2.2.1), we compared the frequency and distribution of single nucleotide variants across UTR pG4 sequences against non-pG4 forming sequences of UTRs. To ensure that only noncoding regions of UTRs were assessed, we removed fractions of UTRs overlapping with annotated coding sequence (CDS) from consideration. To assess evolutionary constraint over UTR pG4 sequences, we compared the allele frequencies of single nucleotide variants (SNVs) affecting these regions, under the expectation that evolution removes deleterious variation from the population. Because of this negative (purifying) selection, we hypothesized that allele frequencies for variants affecting UTR pG4 sequences should be skewed towards more rare variation compared to non-pG4 UTR variants.

Allele frequencies throughout the genome are often affected both by local sequence context, which influences the mutability of a base at a given position, and nearby constrained functional elements that are under linked selection. To control for these confounding factors, we selected a subset of transcripts whose estimated levels of constraint matched our set of UTR pG4-containing mRNAs. Transcript-level constraint was matched using the “observed vs. expected” (o/e) metric, representing the number of loss-of-function mutations occuring within protein-coding sequences of each transcript as reported by gnomAD (suppl. Figure 1 and **Methods**). We used these constraint-matched transcripts to assess allele frequencies across non-pG4 forming UTRs to control for the effects of nearby constrained genes. To additionally account for differences in individual nucleotide mutation biases, we also compared only variants in guanines within pG4 G-tracts against non-pG4 G-tracts, found within 5’ and 3’ UTRs (see **Methods** for details). **Figure 1b** shows that even after controlling for sequence constraint at the transcript level, and only comparing variants affecting guanines, we observe a significant depletion of variants in pG4 and rG4-seq G4s. Specifically, for pG4 sequences with experimental support (rG4 sequences), we found the number of variants per sequenced allele was approximately one-third of that compared to non-pG4 G-tracts in constraint-matched transcripts in the 5’UTR, and 20% lower for the 3’UTR. For pG4 sequences without direct experimental support, G-tract variant frequency differences were similarly reduced (*P*<5e-18 for 5’UTR, <2e-21 for 3’UTR; Fisher’s Exact Test), but of lesser magnitude. This reduction in average allele frequencies for variants within both 5’ and 3’UTR pG4 sequences relative to those not affecting pG4 sequences is consistent with the effects of negative selection.

To provide a complementary measure of sequence constraint, we assessed the extent of polymorphism within UTR pG4 sequences compared to non-pG4 sequences. Under the hypothesis that UTR pG4 sequences are maintained under negative selection, a reduction in the number of polymorphic sites within UTR pG4 sequences is expected. Since nucleotide substitution probabilities are strongly influenced by local sequence context, we applied a background model of neutral evolution based on heptamer mutation rates that has been shown to explain a median of 81% of the variability in substitution probabilities for noncoding regions of the genome^16^. Using this local sequence model, we assessed the number of *observed* versus *expected* polymorphic sites across UTR regions in the European sub-population of the 1000 genomes project, to control for the possible confounding influences of population structure (Phase 1 release – see methods). To simultaneously test the hypothesis that G-tracts within pG4 sequences exhibit reduced observed / expected substitution ratios due to their involvement in secondary structure formation, we further partitioned UTR pG4 sequences by G-tract forming sequences, and intervening “gap” sequences (**Figure 1c**).

Using substitution probabilities from the local sequence context model, we evaluated the difference in the number of expected versus observed substitutions across UTR pG4 sequences compared to constraint-matched non-pG4 forming UTRs (**Figure 1c**). Again, to control for the possible confounding effects of linked selection driven by nearby constrained coding elements, or differences in sequencing depth across the 1000 Genomes project, we produced an empirical distribution for observed vs. expected substitutions in constraint-matched 5’ and 3’UTR sequences by sampling 5000, 25-nucleotide long subsequences from each UTR, over 10,000 iterations using the same heptamer-based substitution model (Suppl. Figure 2). Consistent with the observed reduction in variant frequencies across UTR pG4s, we find a significant reduction in the number of observed versus expected polymorphic sites within UTR pG4 sequences compared to non-pG4 forming regions of the UTR. Specifically, relative substitution rates in 5’ and 3’ UTR G-tracts are approximately reduced by 20% compared to non-pG4 regions of constraint-matched UTRs (permuted *P*<10^−4^, for 5’ and 3’UTR pG4 and rG4) – **Figure 1c**. Surprisingly, 5’UTR pG4 “gap” sequences exhibit a similar reduction in the observed versus expected substitution rates compared to that expected for 5’UTR G-tracts (permuted *P*<10^−4^), suggesting that the “gap” sequences linking 5’UTR pG4 G-tracts may serve additional roles in G-quadruplex stabilization, or have separate biological functions, including possibly serving as a substrate for protein binding. Relative substitution rates in 3’UTR gap sequences, in contrast, are not significantly different from the background 3’UTR estimate, consistent with a pattern of selective pressure in 3’UTR pG4 sequences that acts primarily to maintain the capacity for secondary structure formation.

Taken together, the reduction in allele frequencies, and in the number of polymorphic sites within UTR pG4 sequences suggest that UTR pG4 sequences are under heightened selective pressures compared to non-pG4 forming UTR regions. To place the degree of selection on UTR pG4s in context, we applied the mutability-adjusted proportion of singletons (MAPS) metric^13^, which measures the strength of negative selection acting on a particular class of variants. Within the canonical pG4 motif, we predicted that variants affecting the central guanine of each G-tract should be most constrained, since biophysical studies of G4 stability have shown that mutations affecting the central tetrad (2nd guanine of each trinucleotide guanine repeat) are most detrimental to secondary structure stability^17^. To remove the ambiguity of which specific guanines are involved in secondary structure formation when more than three guanines form a pG4 G-tract, we focused only on single nucleotide variants within trinucleotide G-tracts (n = 3137). By examining variation across each pG4 G-tract, we found central guanine positions within UTR pG4 G-tracts are consistently enriched for singletons (only one sequenced variant in gnomAD) compared to non-pG4 UTR variants, which exhibit similar degrees of constraint as synonymous coding variants (**Figure 1d** – permuted *P*<10^−4^). Thus, there is an excess of rare variation within pG4 G-tracts, in particular for guanines that are most important for maintaining G4 secondary structure.

The reduction in variation affecting the central guanine was observed in the top one-third of most constrained genes as measured by the gnomAD o/e metric (**Figure 1d**, “LOEUF genes”); and, notably, is comparable to that of missense variants for the subset of pG4-central position variants in ClinVar disease-associated genes (**Figure 1d**, “disease genes”). Surprisingly, the most proximal and distal 5’ and 3’ guanine of each trinucleotide pG4 G-tract demonstrated less consistent enrichment of singleton variants within gnomAD across gene classes, suggesting that these positions are under less negative selection compared to central positions. It is possible that mutations at these positions can preserve the potential for RNA to form non-canonical G4 “2-quartets”. Indeed, these G4 “2-quartets” have been estimated to account for 1/4 to 2/3 of all RNA G4 structures observed by transcriptome-wide rG4-seq^15^.

Taken together, our data shows that UTR pG4 regions exhibit enrichment for rare variation, and a depletion in the number of polymorphic sites compared to expectation based on substitution probabilities derived from sequence context alone. These findings demonstrate that UTR pG4 sequences are under heightened evolutionary constraints compared to non-pG4 sequences within UTRs, and implies that variations within UTR pG4 sequences may have biological consequences for human health and disease.

### Most pG4 motifs within untranslated regions of mRNA are isoform-specific

Several known functional UTR elements, including upstream start codons, Open Reading Frames (uORFs), AU-rich elements, and microRNA binding sites, have been found to be frequently included in alternative 5’ or 3’ UTR isoforms of the same gene^18,19^. Alternative UTR inclusion or exclusion is hypothesized to significantly diversify the number of possible post-transcriptional regulatory interactions for a given gene. Given the observed constraint over UTR pG4 sequences, we hypothesized that, like other functional UTR elements, UTR pG4 sequences should exhibit patterns of alternative inclusion or exclusion.

To evaluate the significance of alternative UTR pG4 inclusion, we mapped UTR pG4s to protein-coding transcripts for each gene in the Ensemble transcriptome database. A gene was considered to harbor a “constitutive” pG4 sequence when all annotated protein-coding transcript isoforms contained at least one pG4. “Alternative” pG4 genes, in contrast, had at least one protein coding transcript isoform lacking a pG4 sequence. Strikingly, we found that over half of all genes containing at least one annotated protein-coding transcript isoform with UTR pG4-forming sequences also encoded for alternative UTRs which lacked pG4 motifs (2254 genes with 5’UTR pG4 motifs and 1425 genes with 3’UTR pG4 motifs – **Fig. 2a**). This distribution of “alternative” and “constitutive” pG4 genes was found to be highly significant through permutation testing, when pG4 and non-pG4 transcripts were randomly assigned to genes (*P*-value<0.0001 for 5’ and 3’ UTRs). Moreover, when we assessed levels of pG4-isoform versus non-pG4 isoform expression across 45 tissues in GTEx, many tissues appear to express both pG4 and non-pG4 transcripts simultaneously (**Fig. 2b**). Notably, in testis, almost half of genes with the capacity for alternative pG4 inclusion within the 5’ or 3’ UTR simultaneously express both pG4 and non-pG4 isoforms. This is in contrast to Whole Blood, where expression of alternative pG4 genes is more frequently restricted to either the pG4 isoform or non-pG4 isoform. From this data, we conclude that most UTR pG4-encoding genes express alternative isoforms which lack pG4 sequences, and that the simultaneous expression of both pG4-isoforms and non-pG4 isoforms is widespread across multiple tissue contexts.

Next, we performed gene ontology analysis to explore the functional associations of alternative UTR pG4 genes. We find that these genes are frequently involved in dynamic intracellular processes, including signal transduction, cellular responses to stress, and metabolic regulation (**Figure 2c** – suppl. Table 2). In contrast, constitutive pG4 genes showed enrichment for biological processes more associated with the activation of gene expression in discrete temporal stages, including those involved in tissue development, pattern specification, and cellular differentiation (suppl. Table 3). These observations, coupled with our finding that many tissues simultaneously express both pG4 and non-pG4 isoforms of the same gene, suggest that isoform-switching between pG4-containing or non-pG4 transcripts may facilitate dynamic cellular responses to external stimuli. More broadly, our results demonstrate considerable variation in alternative pG4 inclusion within UTRs across multiple tissue contexts, and suggest that the relative abundance of pG4 and non-pG4 UTRs may be dynamically regulated within a given tissue.

**Figure 2:**
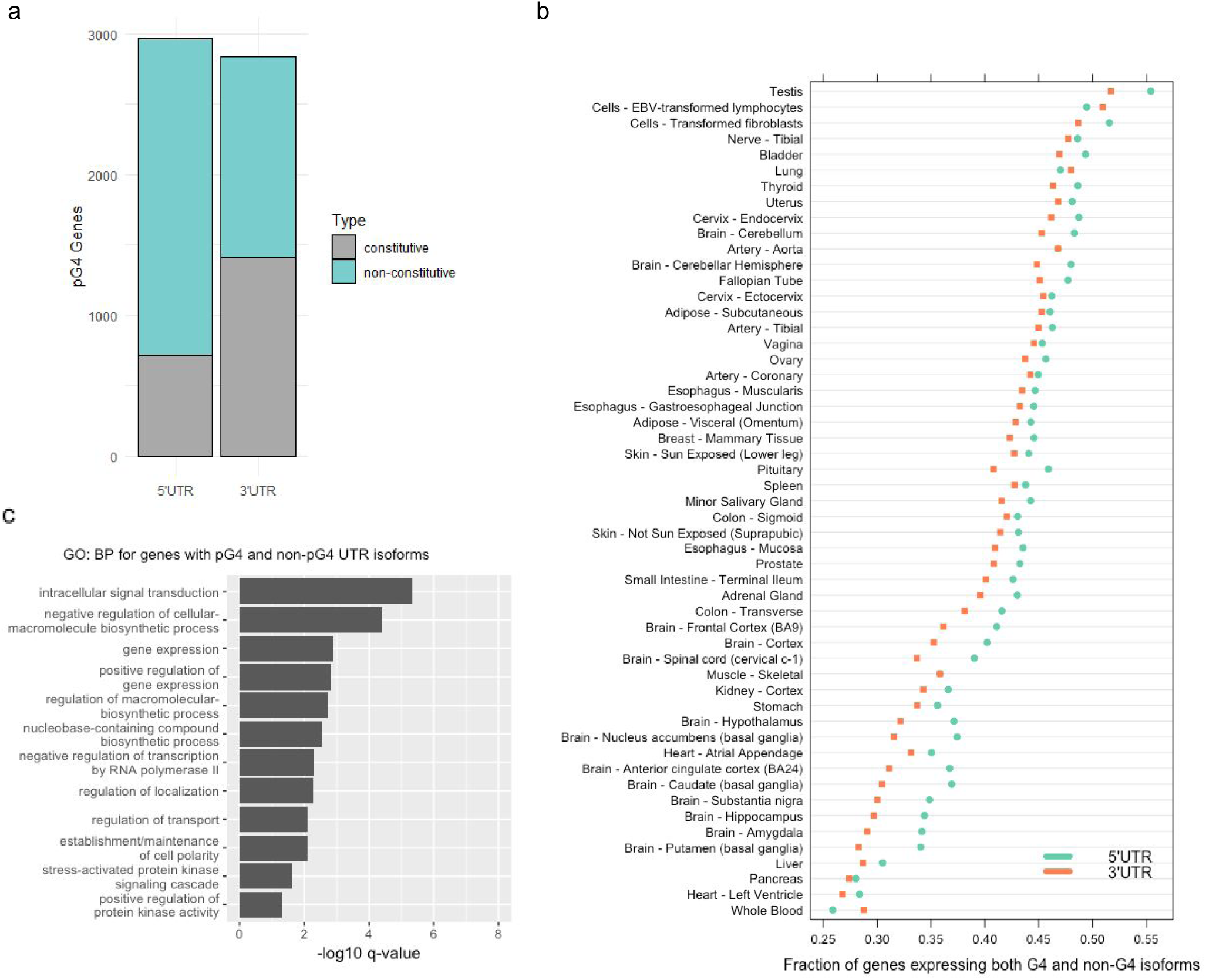
Isoforms with G4 in UTR distribution and usage: **(a)** Most genes producing mRNA transcripts with UTR pG4 sequences also produce alternative isoforms that lack UTR pG4s (non-constitutive). **(b)** For the subset of genes that produce UTRs with alternative pG4 inclusion, both pG4-containing and non-pG4 isoforms are frequently expressed simultaneously. The median TPM measurement for each non-constitutive pG4 encoding gene in each tissue context was aggregated to a composite pG4-”transcript” or non-pG4 “transcript” expression level. Composite pG4 vs. non-pG4 containing transcripts were considered as expressed if their median TPM measurement exceeded 1 TPM for each tissue context considered. The ratio of transcripts expressing both the pG4 isoform, and the non-pG4 isoform was then compared for each non-constitutive pG4 encoding gene, and plotted. **(c)** Overrepresented biological processes for protein-coding genes producing both pG4 and non-pG4 5’ or 3’ UTR isoforms (n = 3148) – see Suppl. Table 1 for full list. GO-term enrichment was performed using PantherDB^49^ and enrichment was determined by meeting av Benjamini-Hochberg adjusted p-value cutoff of 0.05 by Fisher’s Exact Test.

### pG4 motifs in the 5’ and 3’ UTR are enriched for cis-eQTLs

We next evaluated potential regulatory consequences associated with mutations affecting UTR pG4s. We hypothesized that variants occurring within pG4 sequences might also be enriched for variants affecting gene expression. To test this hypothesis, we compared the number of annotated cis-eQTLs versus non-eQTL SNPs identified by the GTEX consortium across pG4 and non-pG4 regions of the UTR. For each gene expressing a transcript isoform containing at least one cis-eQTL, we compared the proportion of non cis-eQTL variants to cis-eQTL variants in pG4 versus non-pG4 UTR sequences. This analysis uncovered significant enrichment for either nominally significant, or lead eQTL variants (lowest P-value variant) in both 5’ and 3’ UTR pG4 sequences compared to non-pG4 regions of UTRs (**Figure 3a** – 5’UTR pG4 Odds Ratio 2.08, 95% CI 1.39 – 3.05, *P*<5×10^−4^, 3’UTR pG4 Odds Ratio 4.96, 95% CI 2.43 – 9.30, *P*<2×10^−5^ for lead cis-eQTLs, 5’UTR pG4 Odds Ratio 1.50, 95% CI 1.22 – 1.83, *P*< 2×10^−4^, 3’UTR pG4 Odds Ratio 2.95, 95% CI 2.33 – 3.73, *P*<2×10^−16^ for nominal cis-eQTLs).

**Figure 3:**
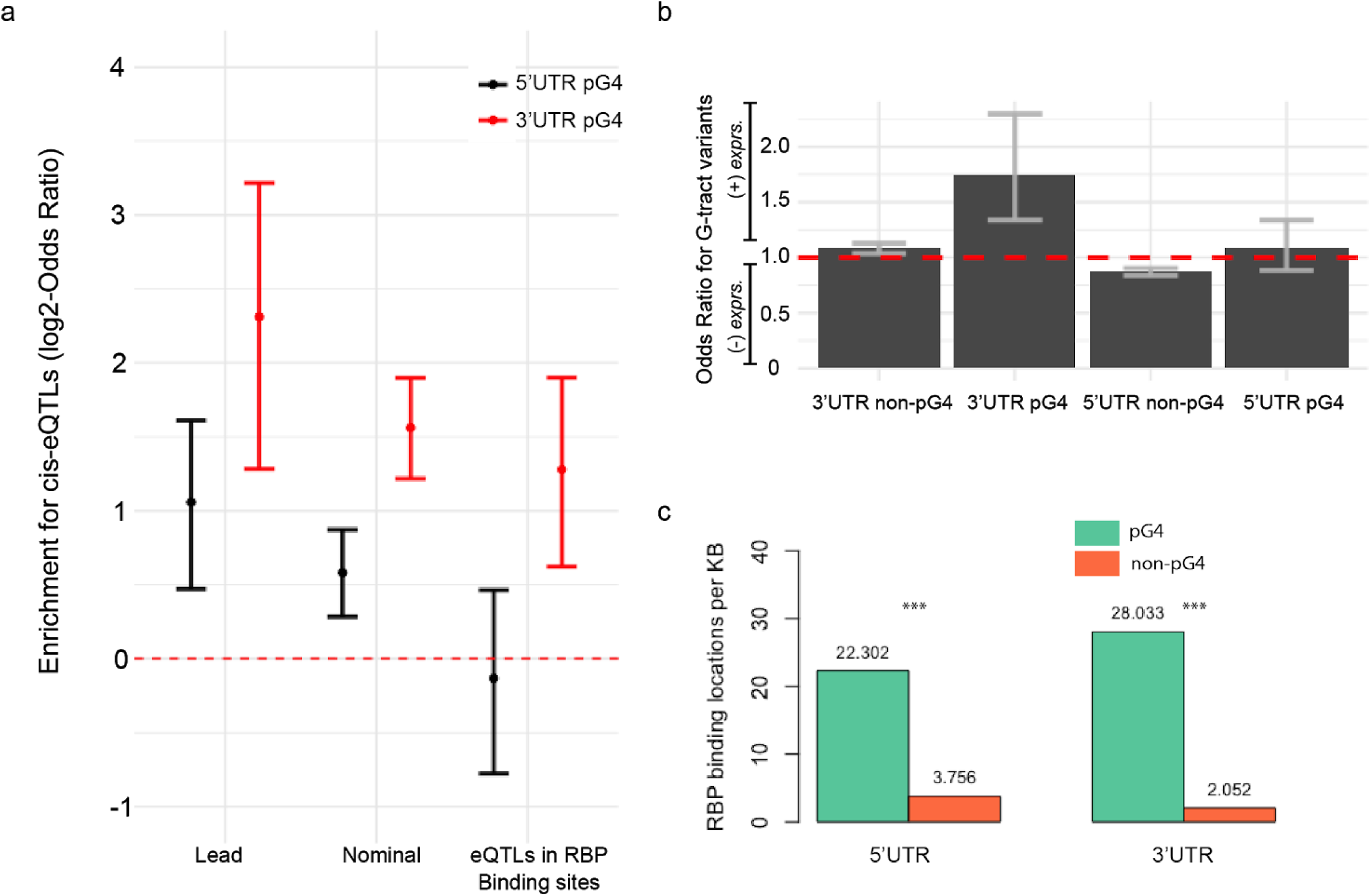
Enrichment for functional associations within UTR pG4 sequences. (a) Lead (n = 256), nominally significant (n = 904), and nominally significant variants in RBP binding sites (n = 288), GTEx v7 expression quantitative trait loci (eQTLs) are overrepresented relative to the number of tested (non-eQTL) SNPs in matched UTR regions (lead = 76,079, nominal = 241,157, RBP overlaps = 28,833, *P*-value<5e-4 for 5’UTR lead, <2e-5 for 3’UTR lead, <2e-4 for 5’UTR nominal, < 2e-16 for 3’UTR nominal, =0.76 for 5’UTR RBP, <2e-4 for 3’UTR RBP, Fisher’s Exact Test). (b) Odds-ratio for a cis-eQTL increasing gene expression across all cis-eQTL-tissue effects (n = 379,441, *P*-value <2e-16, Fisher’s Exact Test), where the variant affects a pG4 G-tract compared to those affecting “gap” sequences. (c) Density of RBP binding sites per kilobase of pG4 sequence compared to non-pG4 regions of the UTR (*P*-value <2e-16, Fisher’s Exact Test).

We next explored the direction of gene expression changes for UTR pG4 cis-eQTLs considering all variant-tissue effects separately for each significant variant-tissue interaction. We hypothesized that variants affecting pG4 G-tracts are more likely to disrupt the structural integrity of the RNA G4s, and thus might influence gene expression differently than variants affecting gap (non-G-tract) sequences within pG4 motifs. Since the magnitude of normalized effect-size estimates in GTEx have no direct biological interpretation, we compared differences in the *direction* of variant effects across pG4 and non-pG4 sequences. As expected, UTR variants in non-pG4 regions of the UTR tend to exhibit similar distributions of positive or negative effects on gene expression, regardless of whether the mutation affected a G-tract, or non-pG4 G-tract nucleotide. This suggests that there is no bias in the effect of variation on gene expression in these regions. Strikingly, we found that mutations pG4 G-tracts in the 3’UTR more often increase mRNA expression compared to those affecting non-G-tract bases (OR 1.75, 95% CI: 1.34 to 2.30, *P*<3.0^−5^) – **Fig 3b**. A similar relationship for the 5’UTR was not observed. Given the known role of the 3’UTR in mediating mRNA stability and degradation, this association may reflect the tendency for 3’UTR G4s to regulate RNA expression levels by facilitating their degradation, thus leading to a tendency for overexpression when G-tract bases are mutated within 3’UTR pG4 sequences.

### RNA-protein binding sites are enriched over UTR pG4 regions

To gain insights into what regulatory mechanisms may be involved in pG4 effects on gene expression we investigated the propensity of protein-binding sites to overlap UTR pG4s. Transcriptome-wide RNA structure mapping studies have suggested that most RNA G4 are unfolded in eukaryotes, but not in prokaryotes, leading to the hypothesis that intracellular factors bind RNA G4s to maintain their unfolded state *in cellulo*^*12*^. We hypothesized that UTR pG4 sequences might be enriched for protein binding interactions. To test this hypothesis, we compared the proportion of UTR pG4 sequences overlapped by RNA-binding protein (RBP) binding sites to non-pG4 forming regions of the UTR, utilizing the expansive catalogue of RBP binding sites published by ENCODE (methods)^20^. This data consists of cross-linking immunoprecipitation sequencing (CLIP-seq) peaks, called from K562 or HepG2 cell lines for over 150 RBPs, containing at least one highly reproducible (IDR = 1000)^21^ binding site within the 5’ or 3’ UTR. Strikingly, we found that, when compared to non-pG4 regions of the UTR, the frequency of overlap between unique (non-overlapping) RBP binding sites and pG4 sequences was almost 6-fold (*P*<2×10^−16^, Chi-square test) higher compared to non-pG4 sequences in the 5’UTR (**Figure 3c**). Enrichment of RBP binding locations over pG4 sequences within the 3’UTR was markedly higher (14-fold, *P*<2×10^−16^, Chi-square test). Given the enrichment within UTR pG4s for cis-eQTLs and protein binding sites, we tested for significant colocalization between these two features within pG4s. Taking the subset of UTR pG4 regions overlapped by any protein binding sites, we examined the density of cis-eQTLs in UTR pG4 regions also overlapping CLIP-seq peaks. When all nominally significant cis-eQTLs are considered, we observe a significant enrichment of cis-eQTLs in the 3’UTR that are also protein binding sites (**Figure 3a**), indicating that variants in 3’UTR pG4 sequences may influence gene expression through changing RNA-protein interactions.

Given the observed association between protein binding sites and pG4 sequences, we next asked whether specific proteins’ binding sites are enriched for pG4s. For each protein, we determined the proportion of protein-specific binding sites containing pG4 sequences, against the total background rate of all CLIP-seq binding sites containing pG4 sequences. To determine significant overrepresentation of pG4 sequences within a given protein’s binding sites, we performed a hypergeometric test against the null hypothesis that there was no overrepresentation of pG4 binding sites within the set of a given protein’s binding sites – **Figure 4a**. This analysis revealed enrichment for proteins that have been implicated in RNA G4 binding (GRSF1, FUS), and those that, to our knowledge, have not previously been associated with RNA G4 structures (PRPF4, GTF2F1, and CSTF2T). GRSF1 is a cytoplasmic protein involved in viral mRNA translation, and has recently been shown to play a role in the degradation of G4-containing RNAs in mitochondria ^22,23^. Other proteins with significant enrichment for pG4 binding include those involved in mitochondrial processes (FASTKD2), transcriptional activation (GTF2F1), mRNA transport (FAM120A), mRNA degradation (XRN2, UPF1), in addition to several proteins implicated in RNA polyadenylation and splicing (CSTF2T, PRPF4, RBFOX2), and surprisingly, micro-RNA (miRNA) biogenesis (DCGR8, DROSHA). Interestingly our analysis reveals little bias for binding the 5’UTR versus the 3’UTR for most pG4-enriched protein binding interactions, with 14 out of 20 proteins’ binding peaks showing enrichment for overlap over 5’ and 3’ pG4 sequences in HepG2, and 17 out of 25 for K562 (Suppl. Table 4). A possible explanation for this is that the RNA G4 structure itself is a more important factor for determining G4-protein binding interactions rather than 5’ or 3’ UTR sequence context alone. Additionally, we find that 12 out of 33 proteins significantly enriched for pG4 peaks in HepG2 are replicated in K562 cells, suggesting that pG4-binding proteins can be shared across cellular contexts. Taken together, our data provides evidence that UTR G4 sequences may exert their functional role by providing molecular signals which facilitate specific RNA-protein binding interactions either directly with RNA binding proteins, or perhaps through interactions that involve protein complexes.

**Figure 4:**
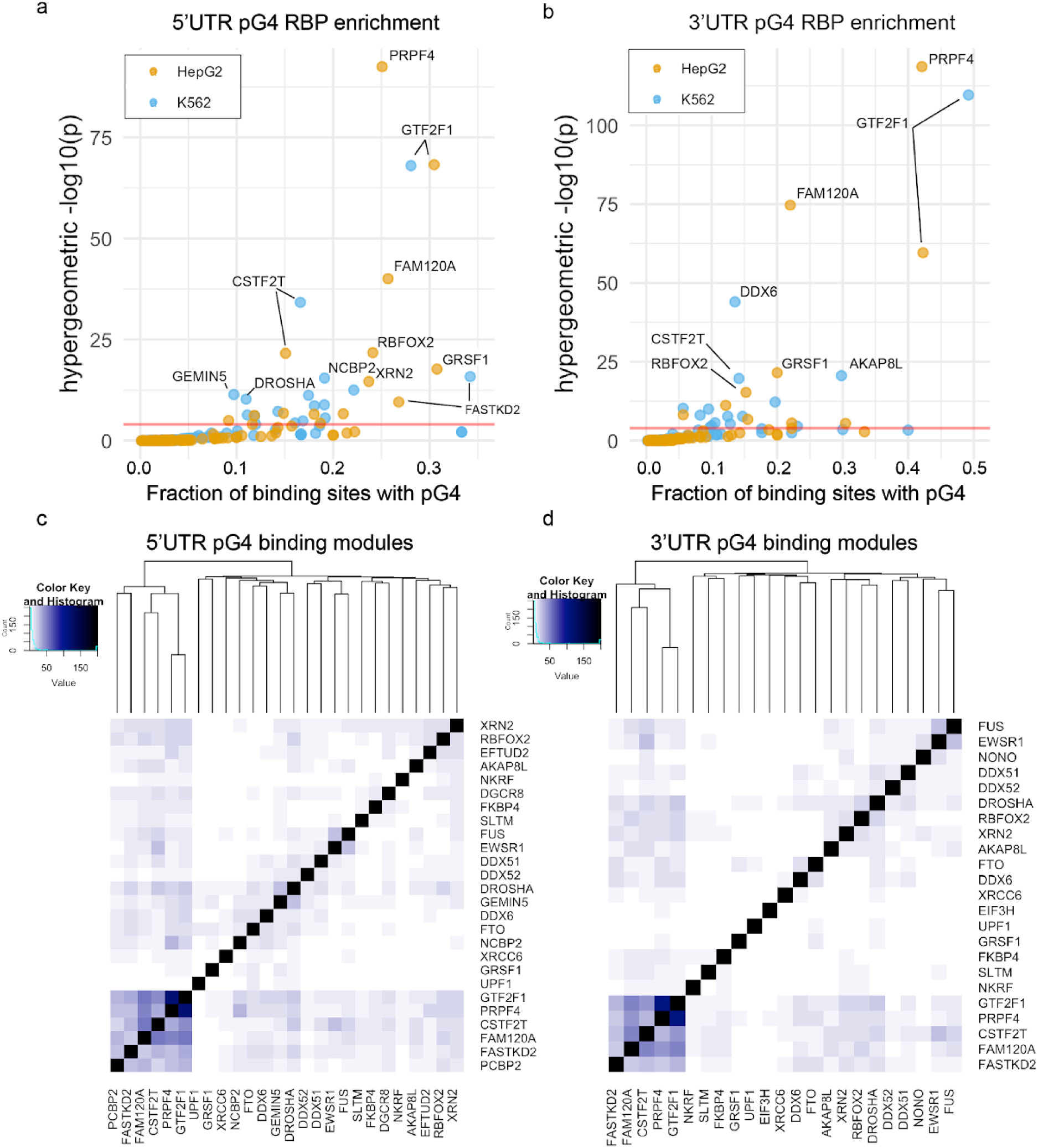
Enrichment of specific protein-pG4 binding sites using CLIP-seq data from ENCODE. (a-b) Enrichment of specific proteins over pG4 binding sites within the 5’UTR (left) and 3’UTR (right) – red line corresponds to p = 0.0001. (c-d) Heatmaps depicting the significance of overlap (hypergeometric -log *P*-value) in pG4 gene targets for proteins found to bind pG4 sequences preferentially. For the subset of proteins that bind UTR pG4 sequences, those involved in RNA processing appear to be the most enriched for pG4’s within their UTR binding sites in both K562 and HepG2 cell lines.

Finally, to explore the potential existence of post-transcriptional regulatory networks relying on shared RNA G4-protein interactions, we tested for a significant overlap in pG4 containing transcripts targeted by each protein enriched for pG4 binding interactions. Taking the set of 31 proteins with significant overrepresentation for pG4 binding (Bonferroni-corrected *P*<0.001) and at least 20 unique pG4 binding sites in HepG2 or K562, we assessed overlaps between the various proteins’ pG4 gene targets. The significance of these target overlaps was tested using a hypergeometric test and visualized using a heat map (**Figure 4c-d**). We did not find high overlap of targets in helicases that have been hypothesized to bind RNA G4s frequently, such as DDX6, DDX51, and DDX52. In sharp contrast, we find a subset of G4-binding proteins sharing a significant degree of overlap in G4-gene targets, including FASTKD2, FAM120A, CSTF2T, PRPF4 and GTF2F1, none of which have been shown to bind RNA G4 structures previously. These data point to possibly novel mechanisms of gene control relying on the shared interactions of these proteins with their respective RNA targets. Indeed, assessing the functional associations of 133 pG4 genes sharing at least 3 out of 5 protein-binding interactions from this enriched module revealed particular association for genes involved in “viral process” (GO:0016032, FDR-adjusted *P*=0.0268, Suppl. Table 4), suggesting that these proteins may be involved in mediating host-viral interactions within the cell.

### pG4s in 3’UTRs of disease-associated genes are enriched for variation

Multiple studies assessing evolutionary constraint in protein-coding regions of the human genome have shown that regions depleted of genetic variation are also enriched for pathogenic variation. Under the principle that purifying selection removes deleterious variants from the genome to produce regions depleted of genetic variation, we should expect that UTR pG4s might also be enriched for pathogenic variation compared to non-pG4 forming regions of the UTR. Since pathogenic variants in ClinVar are overwhelmingly annotated in protein-coding regions of the genome, we were underpowered to test for a direct association between pathogenic variants and UTR pG4 sequences. Instead, we hypothesized that variation is enriched within UTR pG4 sequences of disease-associated genes. To test this hypothesis, we mapped all variants annotated in the most recent release of the ClinVar database (April, 2019) across UTRs, and compared their relative density in pG4 versus non-pG4 sequences in disease-associated genes. We defined the set of disease-associated genes as any gene with at least one variant having an annotation of either “Pathogenic” or “Likely Pathogenic” in ClinVar. We found enrichment for variation in 3’UTR pG4 sequences, annotated 3’UTR rG4-seq G4s, and especially 3’UTR pG4 sequences within annotated RBP binding sites from ENCODE in disease-associated genes compared to non-pG4 forming regions of the 3’UTR – **Figure 5a** (All pG4: OR 1.53, 95% CI 1.28-1.82, *P*<0.00001, rG4-seq pG4: OR 1.16, 95% CI 1.00-1.34, RBP pG4: OR 6.84, 95% CI 4.99-9.16). In contrast, there was no evidence for either enrichment or depletion of pathogenic variants in the 5’UTR, possibly reflecting the generally greater density of other functional elements within 5’UTR sequences. These data suggest pathogenic noncoding variation may be enriched in 3’UTR pG4 regions.

**Figure 5:**
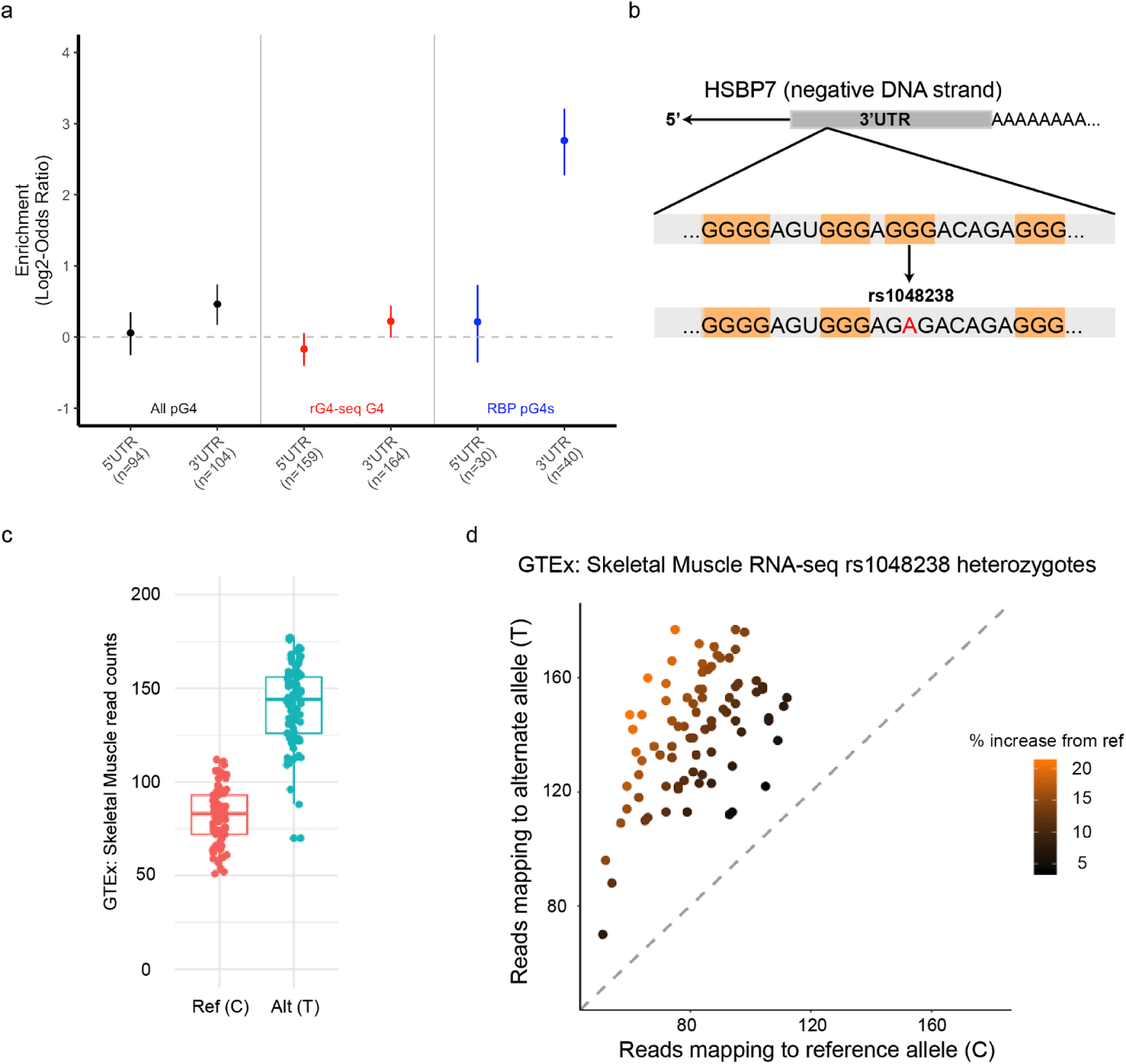
UTR pG4 sequences are enriched for known pathogenic, and putative disease-associated genetic variants. (a) Annotated variants within ClinVar disease-associated genes occur with greater frequency in UTR pG4 sequences compared to non-pG4 UTR regions in the 3’UTR across multiple G4 subsets. (b) rs108348 maps to a 3’UTR pG4 G-tract guanine within the primary HSPB7 transcript, which is encoded on the negative DNA strand. The SNP disrupts the canonical G4 sequence motif by causing a G to A mutation in the RNA transcript. (c-d) WASP-mapping of allele-specific reads in 84 GTEx skeletal muscle samples reveals significant allelic imbalance favoring expression of the alternative allele (*P*-value<1^−100^, likelihood ratio test).

Finally we tested for enrichment of common variants that have been associated with disease phenotypes using annotations available in the NIH GWAS Catalogue. There were not enough GWAS-associated variants within UTR pG4 regions to detect enrichment (8 variants in 5’UTR pG4, 5 in the 3’UTR pG4). However, given the enrichment for cis-eQTLs in 3’UTR pG4, we hypothesized that disruption of UTR pG4 sequences could affect post-transcriptional mechanisms regulating gene expression, providing a potential mechanistic link between GWAS variants and their observed phenotypes. To test this hypothesis, we assessed for evidence of allelic imbalance at rs1048238, a common SNP that has been associated with hypertension in a recent GWAS^24,25^ that affects the central guanine of a pG4 G-tract in the 3’UTR of HSPB7, a chaperone protein that is highly expressed in heart and skeletal muscle (**Figure 5b**). This SNP was selected from the five other 3’UTR GWAS catalogue candidates based on our filtering criteria of having at least 20 heterozygous samples with whole-genome sequencing and matched RNA-seq in disease-relevant tissues from GTEx. Our analysis uncovered a substantial imbalance of reads mapping to the alternative allele of HSPB7 in 84 heterozygous individuals, even after correcting for read-mapping biases using WASP-filtering^26^. Taken together, these results demonstrate that the predicted pG4-disrupting variant is associated with increased expression of the alternative allele at thi s locus (**Figure 5c-d**). This association is consistent with our previous observations from transcriptome-wide mapping of pG4 eQTL showing that 3’UTR pG4 eQTLs tend to increase gene expression (**Figure 3b**); and suggest that the impact of these variants on gene expression are responsible for their respective GWAS associations.

## DISCUSSION

We have applied a deep catalogue human genetic variation to assess evolutionary pressures acting over putative G-quadruplex forming sequences within 5’ and 3’ UTRs. We hypothesized that if these regions are under strong selective pressures compared to the rest of the UTR, they should be depleted of variation. Supporting this hypothesis, we show that variation within UTR pG4 sequences is reduced compared to non-pG4 UTR regions using a local sequence context based substitution model. Moreover, our analysis of positional constraint within the “meta” pG4 motif reveals a particular evolutionary bias for central guanines of each trinucleotide G-tract compared to the terminal positions, and an excess of rare variation affecting these positions implying a similar degree selection as missense variation in disease-associated genes (**Figure 1d**). These findings are consistent with *in vitro* biophysical studies of DNA G-quadruplex stability, which have shown that central position substitutions are most destabilizing, and consequently were predicted to be the most deleterious for native biological functions of G4s^17,27,28^. Interestingly, we find that non-central guanines are less consistently constrained compared to central positions – possibly because mutations at these positions may preserve the potential for RNA to form non-canonical G4 “2-quartets”. Indeed, these G4 “2-quartets” have been estimated to account for 1/4 to 2/3 of all RNA G4 structures observed by transcriptome-wide rG4-seq in HeLa cells^15^.

Our transcriptome-wide pG4 mapping approach also uncovered widespread alternative pG4 inclusion within alternative 5’ or 3’ UTR isoforms (**Figure 2c**). Indeed, alternative 5’ and 3’ UTR usage is a commonly proposed post-transcriptional regulatory mechanism^18,19^, and it is possible that adjusting the inclusion / exclusion of G4 structures specifically contributes to UTR based regulatory mechanisms. For example, global 3’UTR shortening has been observed in cellular responses to arsenic stress, and genes with shortened 3’UTRs are more able to maintain post-stress transcript abundance^29^. Furthermore, we find that genes harboring alternative pG4 UTRs are enriched for biological functions related to intracellular signaling and response to stress stimuli, and show that expression of alternative pG4 containing UTRs is widespread across multiple tissue contexts.

Given the increased sequence constraint over UTR pG4 sequences, we further hypothesized that these regions are enriched for functional associations. GTEx has reported that cis-eQTLs are found more frequently within functionally significant features of the genome, including open chromatin regions and transcription start sites^30^. Here we uncover a greater proportion of cis-eQTLs mapping to pG4 regions compared to non-pG4 sequences within both 5’ and 3’ UTRs. The observed enrichment for cis-eQTLs within 5’UTR pG4 sequences is surprising, since structural elements within 5’UTR G4s have primarily been thought to regulate translational efficiency. Our data suggests that 5’UTR G4s may serve an additional, underappreciated role in mediating mRNA abundance.

Consistent with the role of the 3’UTR in mediating mRNA transcript stability, we observe a greater density of cis-eQTLs in 3’UTR G4s compared to 5’UTR G4s. The 3’UTR is known to contain sequence elements that regulate post-transcriptional gene expression through diverse mechanisms, including miRNA binding sites and the usage of alternative cleavage and polyadenylation (APA) sites ^31^. Two distinct mechanisms describing how 3’UTR G4s can influence post-transcriptional gene expression have been proposed: first, it was shown using luciferase-based reporter constructs that 3’UTR G4 sequences can promote the usage of alternative polyadenylation sites to increase mRNA abundance^11^; and second, 3’UTR G4 secondary structures have been hypothesized to restrict the accessibility of miRNA binding sites to prevent RISC-mediated degradation^10^. Here we demonstrate that variants disrupting pG4 sequences within the 3’UTR are associated with both increased and decreased gene expression depending on the specific variant and tissue context. These findings suggest the roles of 3’UTR G4s in mediating RNA abundance are diverse, and likely context specific.

The activity of proteins which bind RNA G4s may provide one possible mechanistic link between variation in UTR G4s and changes in post-transcriptional regulation. Consistent with this notion, we find evidence that 3’UTR G4s may also mediate interactions with proteins that facilitate RNA degradation given the enrichment of proteins involved in RNA degradation pathways (XRN2, UPF1) over pG4 sequences (**Figure 4c-d**). Indeed, our results show G-tract cis-eQTLs within 3’UTR pG4s are biased towards increasing RNA expression compared to those in non-pG4 sequences. This, coupled with our finding that 3’UTR pG4 sequences are highly enriched for protein-binding interactions more generally, suggests a mechanism where G4-disrupting variation in 3’UTRs confers greater resistance to protein-mediated RNA degradation pathways. Indeed, the RNA-binding protein GRSF1 has been shown to bind and promote G4 melting within mitochondrial-transcribed RNAs that, in turn, facilitates their degradasome-mediated decay^23^. In both K562 and HepG2 cells, we observe similar enrichment for GRSF1 binding over 5’ and 3’ UTR pG4 sequences, leading us to speculate that a similar mechanism may exist to melt pG4 sequences, and promote degradation of their parent RNA transcripts in human cells.

Nevertheless, regulation by protein binding can turn operate via several distinct mechanisms that have already been identified in other examples of UTR-based regulation. Different RBPs can bind similar AU rich elements in the 3’ UTR and exert different effects. Binding of tristetraprolin (TTP) leads to exosome complex recruitment and mRNA degradation while binding of HuR stabilizes mRNA with AU-rich 3’ UTR^32^. In the context of G-quadruplexes, several studies have identified proteins which tend to associate preferentially with DNA and RNA G-quadruplexes using proteomics-based approaches^33–35^. Using CLIP-seq data for over 150 proteins published by ENCODE, we uncovered 40 proteins whose binding sites are enriched for pG4 sequences, and identify regulatory modules associating a set of RNA binding proteins, including FAM120A, FASTKD2, and CSTF2T, with pG4 gene targets involved in viral mRNA expression. Indeed, several examples of viral hi-jacking of eukaryotic RBPs have been reported in the literature^36^, and G4-forming sequences have been found to occur commonly in multiple viral genomes^37^. This, coupled with the observation that RNA G4s appear to be universally depleted within prokaryotic transcriptomes^12^, makes it tempting to speculate that viruses may have evolved G4 sequences specifically as a mechanism for co-opting host cell machinery involved in gene expression and RNA regulation.

Previous approaches for identifying RNA G4-protein interactions have relied on affinity purification assays to uncover proteins that directly bind the folded RNA G4 structure. Since our sequence motif-based approach relies on mapping protein binding sites from CLIP-seq experiments, it is possible that a subset of pG4-enriched proteins identified here might interact with both folded and unfolded G4 structures *in cellulo*. This, coupled with the fact that RNA isoform expression can be highly cell-context dependent, may explain why many of the proteins identified by our approach were not previously reported in the literature.

Nevertheless, enrichment for protein-binding within UTR pG4 sequences using CLIP-seq should be interpreted with some caution given the possibility of proteins binding RNA in complexes, and other higher-order interactions. For example, our analysis showed enrichment for RBFOX2 binding over UTR pG4 sequences, however the canonical RBFOX2 binding motif is well-characterized^38–40^, and does not contain G-rich sequences. Consequently, the observed enrichment of RBFOX2 binding with UTR pG4 sequences is possibly due to secondary to higher-order interactions with other proteins, including HNRNPH, which is known to bind G-rich sequences as part of the LASR complex^41,42^. More generally, this may also explain why many proteins we have uncovered here that are putative G4 binders were not reported as direct RNA G4 binding proteins in previous studies. Additionally, some RNA G4 binding proteins that have been characterized in the literature were not significantly enriched for pG4 binding in the ENCODE CLIP-seq data, likely due to their propensity to also bind non-pG4 containing UTR regions. DDX3X, for example, has been shown to bind RNA G4 sequences^33^, yet it was not significantly enriched for G4 binding in our analysis for this reason.

Finally, given the increased sequence constraint, and enrichment for functional associations within UTR pG4 sequences, we hypothesized that UTR pG4 regions might also exhibit enrichment for pathogenic variation. Using the set of disease-associated genes within ClinVar, we uncovered a greater frequency of single nucleotide variation within 3’UTR pG4 sequences versus non-pG4 UTR sequences (**Figure 5a**). These potentially pathogenic noncoding variants are further enriched for within pG4 sequences that overlap RBP binding sites as annotated in ENCODE. Additionally, we identified a common SNP that has been associated with hypertension within the 3’UTR of HSPB7 (**Figure 5b**), a gene that has been associated with numerous cardiac phenotypes in humans and animal models^43–45^, that is also associated with increased expression of the gene (**Figure 5c-d**), suggesting that the disruption of post-transcriptional regulation within the 3’UTR of HSPB7 may contribute to hypertension. Nevertheless, it remains important to note that the precise causal relationship between these variants and their respective disease associations remains to be elucidated.

An additional caveat to our analysis is that G4 variants associated with gene expression may be acting as markers for other nearby linked features that are causing the observed associations. Although we have uncovered evidence that suggests a causal relationship between UTR G4 sequences and gene expression exists, we cannot rule out the possibility that other cis-regulatory elements in linkage with pG4 variants explain the observed association between the identified pG4 cis-eQTL in HSPB7 and gene expression. The association between allele-specific expression in this gene will require further investigation to establish a causal relationship between G4-disruption and effects on post-transcriptional control of gene expression. More broadly, relatively little is known about the functional consequences of point substitutions within RNA G4s on their functional roles *in cellulo*. Understanding the degree to which RNA G4 sequences can tolerate point mutations while maintaining their biological functions will be important towards extrapolating potential relationships between genetic variants that can be found within these regions and associated phenotypic manifestations. Nevertheless, we present evidence that point mutations within both 5’ and 3’UTR pG4 sequences are associated with changes in gene expression, and speculate that these changes in gene expression could provide a mechanistic basis for the association of certain UTR pG4 variants with disease associations derived from GWAS. Indeed, in aggregate, our findings are consistent with the hypothesis that regions of the genome that are intolerant for variation are also enriched for pathogenic variation.

There are three primary limitations to the current study. First, we applied a text-based pattern-matching approach towards identifying regions of putative G-quadruplex formation within RNA UTRs. Although this approach has been commonly employed in previous work^5,15^, there exists a considerable literature regarding possible variations to the canonical G-quadruplex forming sequence and methods that capture more variable motif definitions ^46,47,48^. Given the comparably limited evidence that many of these alternative G-quadruplex sequences form readily *in cellulo* we used a more stringent canonical motif definition. Nonetheless, it is possible that such alternative G-quadruplex sequences have been missed in our analysis. Thus, our assessment of sequence constraint and functional enrichment within UTR G4 forming regions is likely to be conservative and incomplete. Second, we relied on transcript annotations available in Ensembl to map pG4 sequences within UTRs. pG4 sequences within unannotated UTRs were therefore excluded from our analysis, and consequently the prevalence of RNA G4 structures within UTRs may be more common than reported by our estimate. Finally, although we have uncovered evidence suggesting G4 secondary structure formation is selectively constrained, it remains unclear whether these pG4s form secondary structures *in vivo*. It is possible that some pG4 sequences are able to form secondary structures other than G4s, or that the biological function of these pG4 sequences is unrelated to their capacity for secondary structure formation.

Despite the growing wealth of DNA sequencing data available, our capacity to understand the biological consequences of genetic variation is still limited. Most genetic variants occur within noncoding regions of the genome, and their interpretation within the context of human health and disease remains challenging. Here we provide multiple lines of evidence supporting the biological importance of putative secondary structure forming G-quadruplexes within noncoding regions of the genome. Although RNA UTRs represent only a small fraction of the noncoding genome, they are core components involved in mediating post-transcriptional regulation of gene expression. Thus, we hope this work will motivate researchers to consider G-quadruplexes and other RNA elements in UTRs when assessing the possible impact of genetic variations and interpreting their mechanism of action as it relates to human health and disease.

---

**Suppl. Figure 1:**
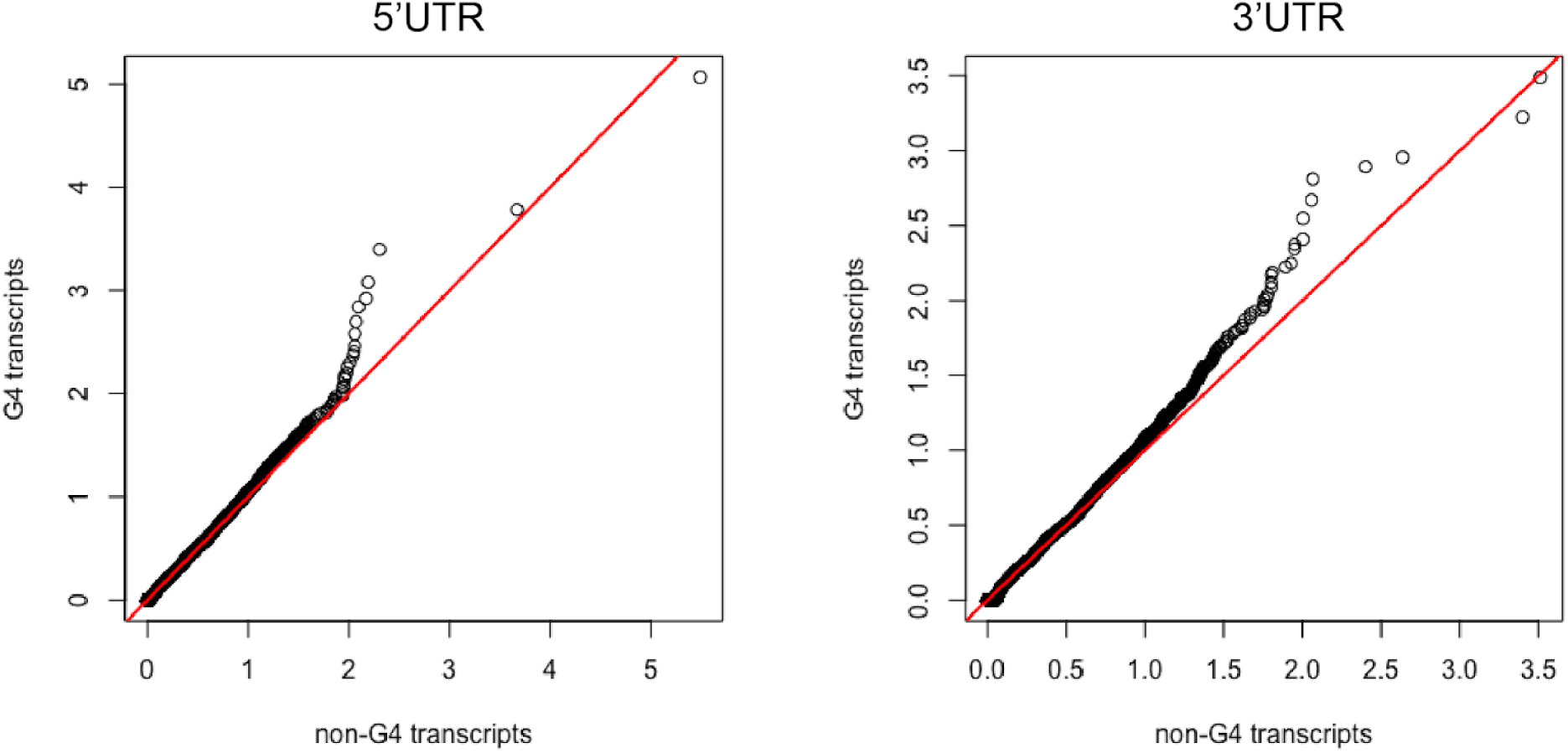
Allele frequencies were compared across constraint-matched transcripts using gnomAD’s observed vs. expected number of loss-of-function mutations (o/e) metric across 5’UTR pG4 and 3’UTR pG4 transcripts.

**Suppl. Figure 2:**
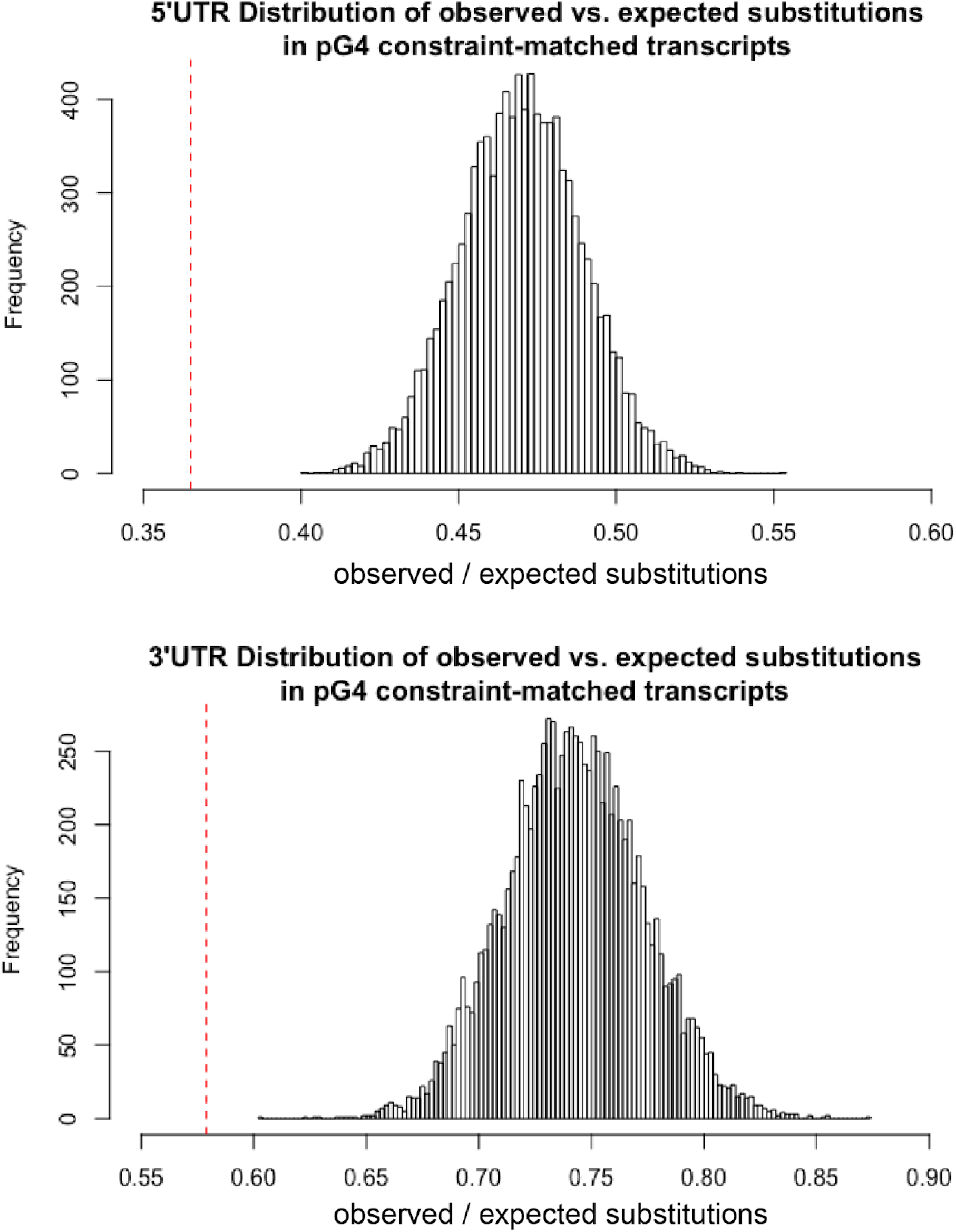
The empirical distribution of observed vs. expected number of substitutions across 10,000 bootstrapped 5’ and 3’ UTR regions in the European subpopulation of the 1000 Genomes Project Phase 1 release. Red dotted line indicates the observed vs. expected ratios estimated by applying the noncoding heptamer mutation model across 5’UTR and 3’UTR pG4 sequences respectively.

## METHODS

### Identification of UTR G-quadruplex Sequences

Annotated UTR sequences and genomic coordinates were downloaded using biomaRt^50,51^ for Ensemble Transcript Database version 75 for all protein-coding transcripts. Putative G-quadruplex forming sequences (pG4) were identified using this set of annotated UTRs by performing a pattern-matching text search to identify regions of UTRs matching the canonical G-quadruplex pattern as a regular expression: (G:3+)-{N:1-7}(3)-(G:3+). Genomic coordinates for pG4 sequences within UTRs were obtained using a custom python script, and cross-referenced with annotated protein-coding regions of the genome from the Ensemble Transcript Database to remove overlaps between annotated UTRs and coding sequences using human genome hg19 coordinates. The identified set of 5’ and 3’ pG4 sequences, and corresponding genomic coordinates were used for all downstream analysis.

A second set of pG4 sequences with evidence of secondary structure formation was defined by overlapping pG4 motifs identified transcriptome-wide with published *in vitro* rG4-seq annotations by using the K+PDS conditions ^15^. Only the subset of rG4-seq G4s matching the canonical pG4 motif were used in this analysis.

### Assessment of Variant Allele Frequencies across UTR pG4 and non-pG4 regions using gnomAD

Variants from gnomAD release 2.2.1 were obtained from (URL here: https://gnomad.broadinstitute.org/downloads) and filtered to exclude those marked with segmental duplication, low complexity regions (LCR), and decoy flags, in addition to those variants whose “True Positive” probability as determined by a random forest model trained in gnomAD did not exceed 40%. As an additional requirement, only those variants where the total observed allele number was at least 80% of the maximum number of sequenced alleles was considered to control for differences in sequencing depth in the gnomAD WGS dataset. The remaining set of high-confidence variants was overlapped with genomic coordinates for UTR pG4, non-pG4, and CDS regions, using bedtools2 (version 2.27.1) intersect with the -u and -b flags.

The transcript constraint-table from gnomAD release 2.2.1 (*URL https://storage.googleapis.com/gnomad-public/release/2.1.1/constraint/gnomad.v2.1.1.lof_metrics.by_transcript.txt.bgz*) was used to randomly select a matching set of transcript-level constraint-matched non-pG4 UTR sequences based on the gnomAD observed / expected metric for the 5’ UTR and 3’ UTR separately. Specifically, transcript constraint was matched between pG4 and non-pG4 forming sequences using the observed versus expected ratio of loss of function variants metric (o/e) provided for each transcript by gnomAD.

The fraction of variants per sequenced allele across UTR regions were computed as the fraction of the observed allele count versus observed allele number. The distribution of frequencies for variants mapping to each UTR region was extracted from the gnomAD summary variant call files directly. P-values for difference between the expected number of variants per sequenced allele across genomic regions were calculated using a two-sided Fisher exact test. Only variants that did not overlap annotated coding regions of the transcriptome were compared to ensure that UTRs overlapping coding regions of other transcript isoforms were excluded. All statistical tests were conducted for 5’UTR and 3’UTR features separately.

For positional constraint analysis we applied the mutability-adjusted proportion of singletons (MAPS) metric^13^ for each nucleotide position across all trinucleotide G-tracts with G4-forming capacity, as defined by our bioinformatic analysis. To control for ambiguity regarding which specific guanines within each G-tract are involved in pG4 formation for G-tracts having more than 3 guanines, we considered only variants within trinucleotide G-tracts. MAPS values were also determined for the set of variants with a VEP consequence of “missense”, or those variants predicted to cause a loss of function (pLoF) in gnomAD to provide context for the different degrees of purifying selection acting over a set of variants. pLoF variants were defined as those annotated with Ensemble predictions for having a high impact and includes “transcript_ablation”, “splice_acceptor_variant”, “splice_donor_variant”, “stop_gained”, “frameshift_variant”, “stop_lost”, and “start_lost” terms. In our assessment of positional constraint within the “meta”-pG4 sequence consisting of only trinucleotide G-tracts, we calculated MAPS for three categories of pG4 variants: 1) all genes, 2) genes with at least one transcript falling in the upper one-third of transcripts that are most intolerant to loss of function mutations (as determined by the gnomAD o/e metric), and 3) disease-associated genes extracted from ClinVar database (April, 2019 release). *P*-values were obtained by performing 10,000 bootstraps for each set of variants in gnomAD.

### Comparison of observed versus expected substitution frequency using the heptamer-based noncoding substitution model

Posterior substitution probabilities for noncoding regions of the genome based on local heptameric sequence contexts were obtained from a published model^16^ based on the Phase 1 release of the 1000 Genomes Project. Cumulative substitution probabilities for each of the possible mutations within a heptamer context (e.g. A->C, A->T, A->G) were calculated by summing over all nucleotide substitution probabilities for a given heptamer context. To produce a null distribution of observed / expected number of substitutions for non-pG4 regions of the UTR, we randomly sampled 5000, 25 nucleotide UTR regions from constraint-matched transcripts, 10,000 times to generate a null distribution. Specifically, constraint-matched non-pG4 transcripts were divided into heptamers using a sliding window across the entire region, and substitution probabilities based on heptameric context alone was summed for each nucleotide position of each region to estimate the expected substitution frequency across each region of interest. The number of expected substitutions as derived from the heptamer substitution model for a given region was compared to observed substitutions for the European subpopulation within the Phase 1 of the 1000 Genomes Release. We performed comparisons across UTR pG4 G-tracts, pG4 non-G-tract “Gap” sequences, and constraint-matched non-pG4 UTRs for all pG4s, and the subset of rG4-seq supported pG4 motifs. Statistical significance was determined by bootstrap resampling from each region of interest 10,000 times and a P-value was calculated as the proportion of resampled observed / expected ratios exceeding a given threshold.

### Assessing pG4 Isoform Expression Across Tissues in GTEx

The median expression of each annotated RNA transcript (as measured in units of TPMs) in each tissue context was downloaded from GTEx. Median TPMs for each transcript were summed for all pG4- or non-pG4 containing transcript for each pG4 gene, and a threshold of 1 TPM was used to determine expression within a specific tissue context. pG4 transcripts were deemed “constitutive” if only one of the pG4, or non-pG4 transcripts exceeded this threshold, and labeled “alternative” if both the pG4, and non-pG4 transcripts exceeded this threshold. Significance of the distribution of “alternative” versus “constitutive” UTR pG4-encoding genes was assessed by randomly assigning pG4 and non-pG4 transcripts each gene, maintaining the number of transcript isoforms encoded by each gene constant with the condition that each gene should contain at least 1 pG4-encoding transcript. The distribution of the ratio of “alternative” to “constitutive” pG4 genes from the randomly distributed pG4 transcripts was then computed over 10,000 bootstraps to obtain a P-value for the true ratio of “alternative” to “constitutive” UTR xpG4 encoding genes.

### Assessment of cis-eQTL and protein-binding enrichment using GTEx and ENCODE

Significant variant-gene pairs were obtained from GTEx release version 7. Lead cis-eQTL variants for each gene were computed using a custom shell script that selected the variant with the lowest *P*-value for each gene, from the set of all significant variants for each tissue context separately. The set of lead and nominally significant variants was overlapped with UTR pG4 and non-pG4 regions of the UTR, and the number of significant cis-eQTL variants per region was compared to the number of non-significant tested SNPs occupying the same region to determine the proportion of cis-eQTLs compared to non-cis-eQTL SNPs in pG4 regions versus non-pG4 regions. UTRs with cis-eQTLs not associated with the same gene giving rise to the UTR were excluded this analysis. Enrichment of cis-eQTLs was computed using a two-sided Fisher Exact Test.

The direction bias of nominally significant cis-eQTLs within UTR pG4 G-tracts versus non-G-tract variants was computed by binarizing the normalized effect size pre-computed for each QTL by GTEx, and comparing the proportions of QTLs in each feature with either a positive effect, or negative effect on gene expression for each possible cis-eQTL annotation across all tissue contexts combined. Statistical significance was determined by a two-sided Fisher Exact Test.

High-confidence protein-binding sites were obtained from ENCODE CLIP-seq summaries and only peaks called with an Irreproducible Discovery Rate = 1000 were used for downstream enrichment analyses as determined by ENCODE ^21^. Overlapping binding sites for multiple proteins were collapsed into a single protein-binding site, and the density of unique binding sites overlapping UTR pG4 regions compared to non-pG4 regions of the UTR was compared by dividing the number of CLIP-seq peaks overlapping each feature by the total number of nucleotides in each region. Significance was assessed using a chi-square test with 2 degrees of freedom.

Proteins whose binding sites are enriched for pG4 overlaps were computed using a hypergeometric test, taking the set of pG4 containing protein-specific binding sites compared against the background frequency of all observed UTR CLIP-seq peaks containing a pG4 sequence. The significance of pairwise overlaps between protein-gene targets was also computed using a hypergeometric test to assess the degree that one protein’s pG4 binding genes were also targets for another protein.

### Assessment of overlap between pathogenic genes in ClinVar and the NIH-GWAS Catalogue

The April 2019 release of ClinVar was obtained from ftp://ftp.ncbi.nlm.nih.gov/pub/clinvar/. Using these variant annotations, we identified a subset of disease associated genes as any gene with at least one variant having a “Pathogenic” or “Likely Pathogenic” annotation. These genes were used to subset the ClinVar database and single nucleotide variants were overlapped with UTR regions to assess for enrichment in pG4 sequences. Insertions, deletions, and expansions were not considered in this analysis. The number of variants across each region was then divided by the total number of bases in each respective region to estimate of the density of variation in a given region. The odds ratios for the number of single nucleotide variants compared to the number of bases in a given region were then compared using a two-sided Fisher Exact Test. Publicly available phenotype-associated SNPs from were obtained from the NIH-EBI GWAS Catalogue.

### Sequencing alignment, mapping, and allele-specific expression for GWAS-variants

RNA-seq libraries were trimmed using TrimGalore^52^. Reads were aligned to the GRCh37 human genome using STAR (version 2.7.0c) with the WASP-filtering option and matched whole-genome sequencing variant files obtained from GTEx for Skeletal Muscle and Tibial Nerve tissue samples. Reads that did not pass WASP-filtering were removed from the resulting aligned bam files. PCR duplicates were removed using the python script remove_duplicates.py included in the WASP version 0.3.3 pipeline (https://github.com/bmvdgeijn/WASP). Read counts matching the reference and alternate alleles in the resultant WASP-filtered bam files were compiled using bcftools mpileup across UTR pG4 variants. A beta-binomial model was fitted using the *R VGAM* package^53^ for each variant across all heterozygous samples identified using matched whole-genome sequencing from GTEx to estimate the ratio of reference reads to alternate reads. Estimates of statistical significance were obtained by using a likelihood ratio test comparing the log-likelihood of the observed count distribution for each variant using the beta-binomial estimate for /rho versus the null hypothesis of no bias (/rho = 0.5).

### Statistics

Data were analyzed and statistics performed using *R* (version 3.5.0) and *Python* (version 3.7 and 3.6.1). Significant differences are noted by asterisks (***).

